# Cell-matrix adhesion contributes to permeability control in human colon organoids

**DOI:** 10.1101/2021.06.11.448147

**Authors:** James Varani, Shannon D. McClintock, Muhammad N. Aslam

## Abstract

**Background and aims:** Cell-cell adhesion structures (desmosomes and, especially, tight junctions) are known to play important roles in control of transepithelial permeability in the colon. The involvement of cell-matrix interactions in permeability control is less clear. The goals of the present study were to: i) determine if disruption of colon epithelial cell interactions with the extracellular matrix alters permeability control and ii) determine if increasing the elaboration of protein components of cell-matrix adhesion complexes improves permeability control and mitigates the effects of cell-matrix disruption.

**Methods:** Human colon organoids were interrogated for transepithelial electrical resistance (TEER) under control conditions (0.25 mM calcium) and in the presence of Aquamin^®^, a multi-mineral product, at a level providing 1.5 mM calcium. The effects of Aquamin^®^ on cell-matrix adhesion protein expression were determined in a proteomic screen and by Western blotting. In parallel, TEER was assessed in the presence of a function-blocking antibody directed at an epitope in the C-terminal region of laminin α3 chain.

**Results:** Treatment of colon organoids with Aquamin^®^ increased the expression of multiple basement membrane and hemidesmosomal proteins as well as keratin 8 and 18. TEER values were higher in the presence of Aquamin^®^ than they were under control conditions. Anti-laminin antibody reduced TEER values under all conditions but was most effective in the absence of Aquamin^®^, where laminin expression was low and TEER values were lower to begin with.

**Conclusions:** These findings provide evidence that cell-matrix interactions contribute to permeability control in the colon. They suggest that the elaboration of proteins important to cell-matrix interactions can be increased in human colon organoids by exposure to a multi-mineral natural product. Increasing the elaboration of such proteins may help to mitigate the consequences of disrupting cell-matrix interactions on permeability control.

## INTRODUCTION

Functional defects in the gastrointestinal tract barrier have been documented in inflammatory conditions of the bowel, including both ulcerative colitis (UC) and Crohn’s Disease (1-5). Barrier defects have also been described in irritable bowel syndrome (6) and noted in celiac disease (5) and as a consequence of acute bacterial infection (7). Barrier defects have also been seen in obesity related to high-fat and high-sugar diets (8,9) and, thus, may contribute to chronic, systemic inflammation. Finally, gastrointestinal discomfort associated with chronic environmental stress may reflect barrier dysfunction related to corticosteroid release (10). In these situations, inflammatory injury to the intestinal wall contributes to barrier breakdown. At the same time, however, pre-existing barrier defects, leading to permeation of bacteria, bacterial products, food allergens and toxins into the mucosal wall, may promote inflammation in the gastrointestinal tract (4).

Tight junctions are thought to be the epithelial cell surface structures that most directly contribute to permeability control – at least in so far as soluble factors are concerned (11-17). Desmosomes are also important, if for no other reason than that they are largely responsible for tissue cohesion and strength (11-15,18-20). It is difficult to envision effective control of permeability in a mechanically-active tissue such as the colon if tissue cohesion is compromised. In support of this, we recently demonstrated that Aquamin^®^, a calcium- and magnesium-rich, multi-mineral product obtained from marine red algae (21), strongly up-regulated desmosomal proteins and increased the number of desmosomes in human colon tissue in organoid culture but had little effect on tight junctional elements (22-24). In parallel with these desmosomal changes, both permeability control and tissue cohesion were increased (23). The same multi-mineral intervention that increased desmosomes also up-regulated expression of other moieties that contribute to the permeability barrier. Among these were cadherin family members, carcinoembryonic antigen cell adhesion molecules (CEACAM), mucins and trefoils.

In the present study, we have used a proteomic screen to assess the expression of proteins involved in cell-matrix interactions in human colon organoid culture. Studies by other investigators have utilized immunohistochemical methods to show basement membrane defects in UC and Crohn’s disease as well as in other inflammatory conditions of the bowel (25-28).

While these findings suggest a role for cell-basement interactions in barrier function, how these interactions influence gastrointestinal permeability, *per se*, has not been studied. Here it is demonstrated that proteins affecting cell-basement membrane interactions through both focal adhesions and desmosomes are increased in response to Aquamin^®^. Further, it is shown that an antibody to the major cell adhesion domain in the laminin α-chain decreases transepithelial electrical resistance (TEER) – i.e, a surrogate for permeability control – in human colon organoid culture. In contrast, treatment with the same antibody has no apparent effect on tissue cohesion / tissue strength.

## MATERIALS AND METHODS

### Aquamin^®^

This is a calcium-rich, magnesium-rich, trace element-rich multi-mineral product obtained from the skeletal remains of the red marine algae, *Lithothamnion sp* (21) (Marigot Ltd, Cork, Ireland). Aquamin^®^ contains calcium and magnesium in a molar ratio of approximately 12:1 along with measurable levels of 72 other trace minerals (essentially all of the trace elements algae fronds accumulate from the deep ocean water). The same single batch of Aquamin^®^ Soluble that was used in the previous colon organoid studies (22-24) was used for this study. File S1 describes the complete mineral/trace element composition of the multi-mineral product - Aquamin^®^.

### Anti-laminin antibodies and other reagents

The known laminin α chains contain a globular region in the C-terminal end of the molecule made up of 5 modules (referred to as Laminin G domain-like modules). Cell-binding sites are present within these modules (29-31). A mouse monoclonal antibody (IgG1 clone) prepared against an epitope within the G domain of the human laminin α3 chain was used for functional blockade. This antibody (clone #P3H9-2; R&D Systems) has been demonstrated to detect antigen in a variety of epithelia and has been shown to inhibit cell proliferation of both rat and human epithelial cells (32). A control mouse monoclonal IgG1 immunoglobulin was used in parallel with the anti-laminin antibody for comparison. A rabbit polyclonal antibody (Invitrogen; PA5-27271) prepared against a recombinant protein fragment from the human laminin β1 chain was used in western blotting. A monoclonal antibody recognizing a human actin epitope (Cell Signaling Technology; 5125S) was used as control.

### Organoid culture

Colon tissue in organoid culture was available from our previous studies (22-24). The Institutional Review Board at the University of Michigan Medical School (IRBMED) approved the tissue collection and use protocol. Subjects provided written informed consent prior to flexible sigmoidoscopy and biopsy collection. This study was conducted according to the principles stated in the Declaration of Helsinki. For the present work, cryopreserved colon organoid samples were put into culture and expanded over a 3-4 week period. After expansion, organoids were used to assess transepithelial electrical resistance (TEER) (below) or tissue cohesion (below). Additional organoid samples were used for Western blotting. Three media formulations were used in this study. These include:

#### Growth medium

Growth medium consisted of a 50:50 mix of Advanced DMEM (Gibco) and the same base media that had been conditioned by the previous growth of L-cells engineered to provide a source of recombinant Wnt3a, R-spondin-3, and Noggin – referred to as L-WRN conditioned medium (33). The growth medium was supplemented with 100 ng/mL human recombinant epidermal growth factor (EGF) (R&D) as the major growth-supporting peptide and was further supplemented with 10 µM Y27632 (Tocris), 500 nM A83-01 (Tocris), 10µM SB202190 (Sigma), 2.5 µM CHIR99021 (Tocris), 1X B-27 without vitamin A (Invitrogen), 1 mM N-Acetyl-L-cysteine, 10 mM HEPES (Invitrogen), 2 mM Glutamax (Invitrogen), 10 μM SB202190 (Sigma), and 100 μg/mL Primocin (InvivoGen). Since L-WRN medium was supplemented with 20% fetal bovine serum, after 1:1 dilution, the final concentration of the growth medium was 10% serum.

#### Differentiation medium

Differentiation medium consisted of a mix of Advanced DMEM and F12 media. This formulation lacked Wnt3a and R-spondin-3 but was supplemented with EGF (50 ng/mL) along with Gastrin (10 nM, Sigma), Noggin (50 ng/mL, R&D), and Y27632 (2.5 µM, Tocris). AlbuMAX^®^ (Gibco), a lipid-rich Bovine Serum Albumin (BSA), was used as a component of the medium to replace serum. The final calcium concentration in complete differentiation medium was 1.04 mM.

#### Growth medium-KGM Gold mix

KGM Gold is a serum-free, calcium-free medium designed for epithelial cell growth (Lonza). In the growth medium-KGM Gold mix (at a 1:4 dilution), the serum concentration was 2.5% and the calcium concentration was 0.25 mM (and used as a control). When Aquamin^®^ was included, it was added at 0.51 mg/mL; an amount to bring the final calcium concentration to 1.5 mM.

### Assessment of TEER

Organoids were dissociated into small cell aggregates (less than 40µm in size) and plated onto collagen IV (Sigma)-coated transwells (0.4 µm pore size, 0.33cm^2^, PET, Costar) at 200,000 individual organoids per well in growth medium as in Zou et al. (34). After seeding in growth medium, organoids were allowed to attach to the TEER membrane and incubated without further treatment for one day. Then growth medium was replaced with either differentiation medium or with the growth medium-KGM Gold mix with or without Aquamin^®^.

The function-blocking anti-laminin antibody was included at the start of the treatment period at 25 μg/mL in each of the culture media. Fresh culture medium and antibody was provided every two days during the assay period. A control mouse IgG was used at the same concentration for comparison. TEER values were determined daily beginning on day-2 using an epithelial volt/ohm meter (EVOM2, World Precision Instruments) and STX2 series chopstick electrodes as described previously (23).

### Histochemical staining and light microscopy

At the completion of TEER measurements on day-2 and day-5), membranes with cells still attached were prepared for light microscopy. The membranes were fixed for 1 hour in 10% buffered formalin. Following this, membranes were paraffin-embedded, sectioned and stained with hematoxylin and eosin. The stained specimens were visualized by light microscopy. Slides were digitally scanned using the Aperio AT2 brightfield whole slide scanner (Leica Biosystems) at a resolution of 0.5 µm per pixel with 20X objective.

### Western blotting

At the completion of the TEER measurements, organoids were harvested for protein. Briefly, wells were washed gently with PBS, then harvested using 0.4% EDTA to dissolve and remove matrigel. The remaining pellet containing organoids was then washed 3 times with PBS and subjected to extraction using RIPA buffer (89901; Thermo Scientific). Organoids were lysed by repetitive pipetting in the buffer, followed by incubation for 10 minutes on ice. Non-soluble cellular debris was removed by centrifugation at 14,000g for 10 minutes and protein was quantified using a BCA assay (23227; Pierce). Samples were heated for 10 minutes at 70°C in NuPage LDS sample buffer and then run on 3-8% Tris-Acetate gels using NuPage MOPS running buffer under reducing conditions. Proteins were then transferred onto nitrocellulose membranes, blocked with 5% non-fat dry milk and probed with the primary and appropriate secondary antibodies. Secondary antibodies were used at 1:5000 for all membranes. β-actin was used as a loading control in each assay. SuperSignal WestPico Plus (34577; Thermo Scientific) detection reagent was used and bands were visualized using a BioRad ChemiDoc XRS+ Molecular Imager. Relative band density was determined using Image J gel analysis tools.

### Confocal fluorescence microscopy

At the completion of the TEER assay, membranes were prepared for confocal fluorescence microscopy and stained with an antibody to occludin for the purpose of visualizing the cell layer. The membranes with cells still attached were fixed for 15 minutes at -20°C in methanol. They were then washed three times in PBS before blocking in 3% BSA (A8806; Sigma) in PBS for 1 hour. Following this, membranes were stained with an antibody to occludin (331594; Invitrogen) 1:400 for 1 hour in 1% BSA in PBS. Stained membranes were rinsed 3 times (5 minutes each) in PBS, stained with DAPI for 5 minutes to identify nuclei and washed an additional 3 times with PBS. Finally, the membranes were gently cut from the transwell insert and mounted apical side up on Superfrost Plus glass slides (Fisher Scientific, Pittsburgh, PA) with Prolong Gold (P36930; Life Technologies Molecular Probes). The stained specimens were visualized and imaged with a Leica Inverted SP5X Confocal Microscope System (University of Michigan Medical School Biomedical Research Core Facility).

### Organoid cohesion assay

Organoid cohesion was assessed as described previously (23). Briefly, after establishment and culture expansion, organoids were incubated in growth medium-KGM Gold with or without the same anti-laminin antibody (25 μg/mL) as described above. Treatment was for seven days with fresh medium and antibody added at days 2 and 4. At the end of the incubation period, phase-contrast microscopy (Hoffman Modulation Contrast - Olympus IX70 with a DP71 digital camera) was used to capture images in order to measure the size of multiple individual organoids (53-104 individual organoids per condition). Then organoids were then separated from the Matrigel and fragmented with mechanical force alone by pipetting the entire pellet 30x through an uncut 200 microliter pipet tip. After washing 3x in PBS, organoids were re-cultured in fresh Matrigel. One day after establishment, multiple colonoids (colon organoids) were again examined under phase-contrast microscopy and sized. For both pre-harvest and post-harvest samples, phase-contrast images were analyzed using area measurements in Adobe Photoshop (CC version 19.1.5). Average organoid size-reduction (fold-change before and after harvest and mechanical fragmentation) was determined by dividing the average post-harvest surface area by the average pre-harvest area.

### Differential proteomic analysis

Proteomic assessment was conducted at the Proteomics Resource Facility (PRF) in the Department of Pathology at the University of Michigan using mass spectrometry (MS)-based tandem mass tag (TMT) analysis (ThermoFisher Scientific). The complete details for the experimental conditions, protocols and analysis methodology can be found in previously published reports (22,24). Briefly, fifty micrograms of organoid protein from each condition were digested with trypsin and individually labeled with isobaric mass tags. Labeled peptides were fractionated using 2D-LC (basic pH reverse phase separation followed by acidic pH reverse-phase) and analyzed on a high-resolution, tribrid mass spectrometer (Orbitrap Fusion Tribrid, ThermoFisher Scientific) using conditions optimized in the PRF. MultiNotch MS3 was employed to obtain accurate quantitation of the identified proteins/peptides. Data analysis involved peptide filtering to retain only those that passed ≤2% false discovery rate (FDR) threshold of detection. Quantitation was performed using high-quality MS3 spectra. Differential protein expression values (fold-change) for proteins of interest in each treatment group were compared to protein values of the respective control group. Proteins were identified using Universal Protein Resource (Uniprot) databases (Uniprot.org). Reactome version 75 – a pathway analysis database was used to recognize associated pathways for species “Homo sapiens” (reactome.org). For the purpose of the present study, we accessed two existing data sets – one consisting of colon organoid tissue from four healthy subjects and the other consisting of colon organoid tissue from three ulcerative colitis patients in remission. In all cases, organoids grown in the growth medium-KGM Gold mix were compared to organoids grown in the same medium supplemented with Aquamin^®^ at levels providing 1.5 to 3.0 mM calcium. For comparison purposes, the data presented here include only the maximum response. The complete proteomics data sets are available at the ProteomeXchange Consortium via the PRIDE partner repository with the dataset identifier PXD020244 for UC derived colon organoids (and identifier pending for normal colon organoids).

### Statistical analysis

Means and standard deviations were obtained for discrete values in each assay (TEER, cohesion, and individual proteins from proteomics). Data generated in this way were analyzed by ANOVA followed by paired t-test (two-tailed) for comparison using GraphPad Prism version 8.3.

## RESULTS

### Anti-laminin antibody reduces TEER values in organoids maintained in differentiation medium

In the first series of studies, human colon organoids were plated on transwell membranes in growth medium. One day later, differentiation medium replaced growth medium and electrical resistance was assessed over subsequent days. Results are shown in Fig 1. Fig 1A demonstrates that under conditions optimized to promote electrical resistance (i.e., in differentiation medium), TEER values were low during the first two days after treatment, rose precipitously between days-3 and day-5, and fell thereafter. A combination of antibody to occludin (tight junctional protein) and DAPI (nuclear stain) was used to illuminate organoids on the membranes at day-2 and day-5. As shown in the inserts in Fig 1A, an intact cell-cell border could be seen in organoids by day-2. However, cell outgrowth from the organoids did not completely cover the membrane surface at this time (accounting for the lack of electrical resistance). Coverage of the membrane surface was complete by day-5.

**Fig 1.**
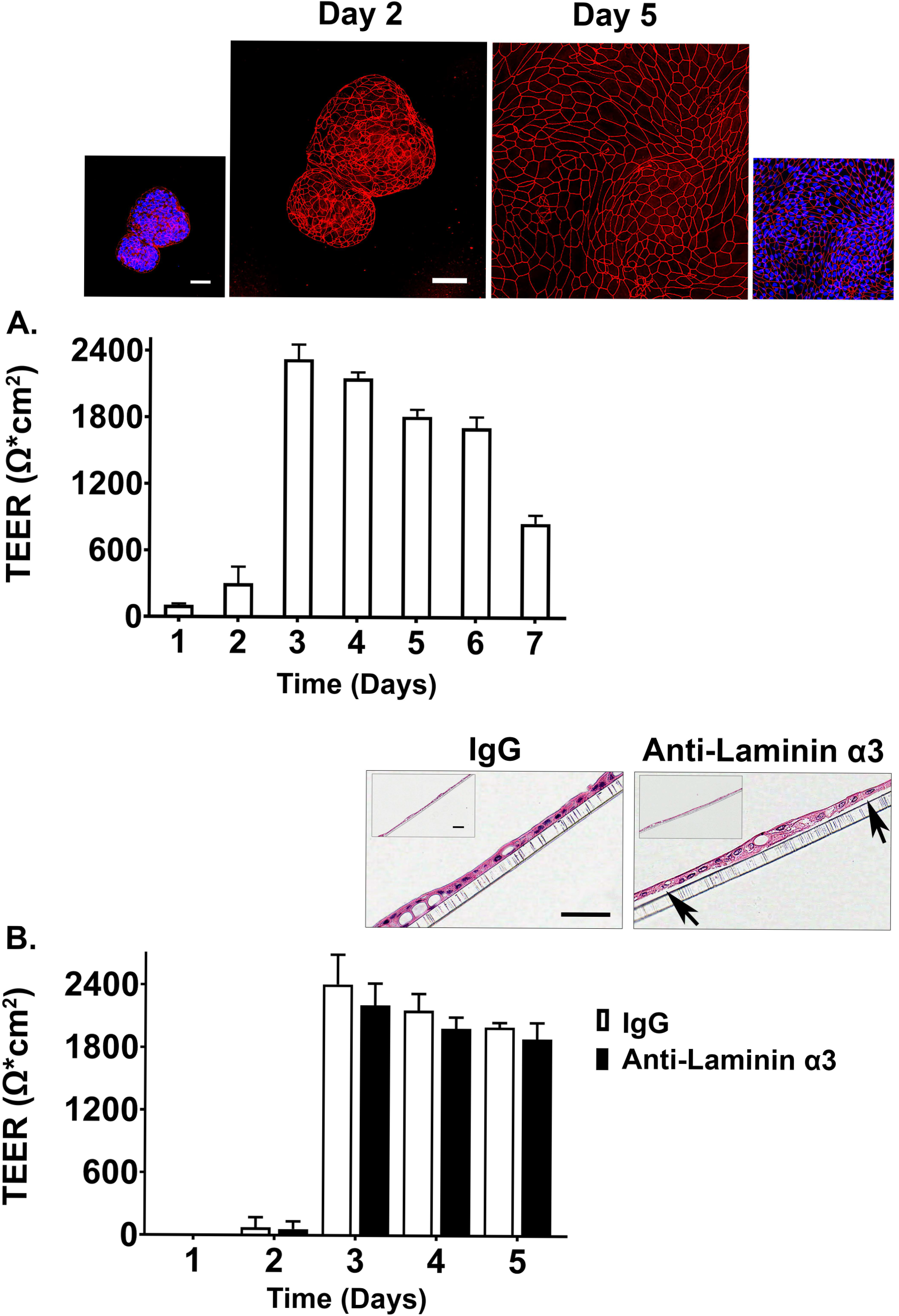
Transepithelial electrical resistance in differentiation medium. **A**: Time-dependent changes in TEER values. Values shown are means and standard deviations based on four separate experiments with 4 samples (individual membranes) per data point at each time-point in each experiment. <u>Insert</u>: Confocal fluorescent microscopic (max-projected) images of membranes stained after the day-2 and day-5 readings with antibody to occludin and with DAPI. Scale bar = 50 µm. **B**: Effects of anti-laminin antibody on TEER values. Values shown are means and standard deviations based on two separate experiments with 4 samples (individual membranes) per data point at each time-point in each experiment. <u>Insert</u>: hematoxylin and eosin-stained images of TEER membranes from IgG-treated and anti-laminin-treated wells. Arrows in the anti-laminin-treated image show areas where cell detachment from the underlying TEER membrane was visible. Scale bar = 100 µm (small) and 50 µm (Large).

Fig 1B demonstrates the effects of a function-blocking antibody to the α3 chain of laminin on electrical resistance in differentiation medium. A modest decrease in TEER values was observed at days-3, -4 and -5 (9-17% decrease). When values from the three time-points were averaged together, the decrease observed with the anti-laminin antibody compared to the IgG control reached a statistically significant level (mean decrease = 11±3%; p<0.01 by 2-tailed t-test).

At the completion of TEER assessment, membranes with cells still attached were fixed in 10% buffered formalin, stained with hematoxylin and eosin and examined at the light microscopic level (Fig 1B insert). It can be seen that under control conditions (IgG-treated cells) or in the same medium with anti-laminin antibody, most of the membrane surface was covered with a complete monolayer of cells. However, in the presence of the anti-laminin antibody, focal areas where cells had detached from the underlying substrate could be observed. In these areas, cell-cell attachments remained intact such the structure had the appearance of a tiny “blister.”

### Aquamin^®^ up-regulates basement membrane components, proteins associated with hemidesmosome formation and keratins

Fig 2A presents data from the proteomic assessment based on merged data from healthy subjects (n=4) and subjects with UC (n=3). In both data sets (assessed independently), strong up-regulation of several laminin chains (α1, β1, β2 and γ1) (components of laminin 111 and 121) along with nidogen-1 and the basement membrane-specific heparin sulfate proteoglycan (HSPG-2, perlecan) was seen. Also detected in the proteomic analysis (Fig 2A) were laminin α3, β3 and γ2 chains (components of laminin 332 or laminin-5 in the older terminology). This laminin isoform is a major component of hemidesmosomes (35,36) and is considered as a skin-specific laminin isoform (37). While these laminin chains did not demonstrate an increase in response to Aquamin^®^ in colon organoids, several other hemidesmosomal proteins were detected, and a subset of these (plectin, desmoplakin and epiplakin) were increased by Aquamin^®^ as compared to control. The plakins are critical linkers between laminin in the hemidesmosomes and intermediate filaments (38,39).

**Fig 2.**
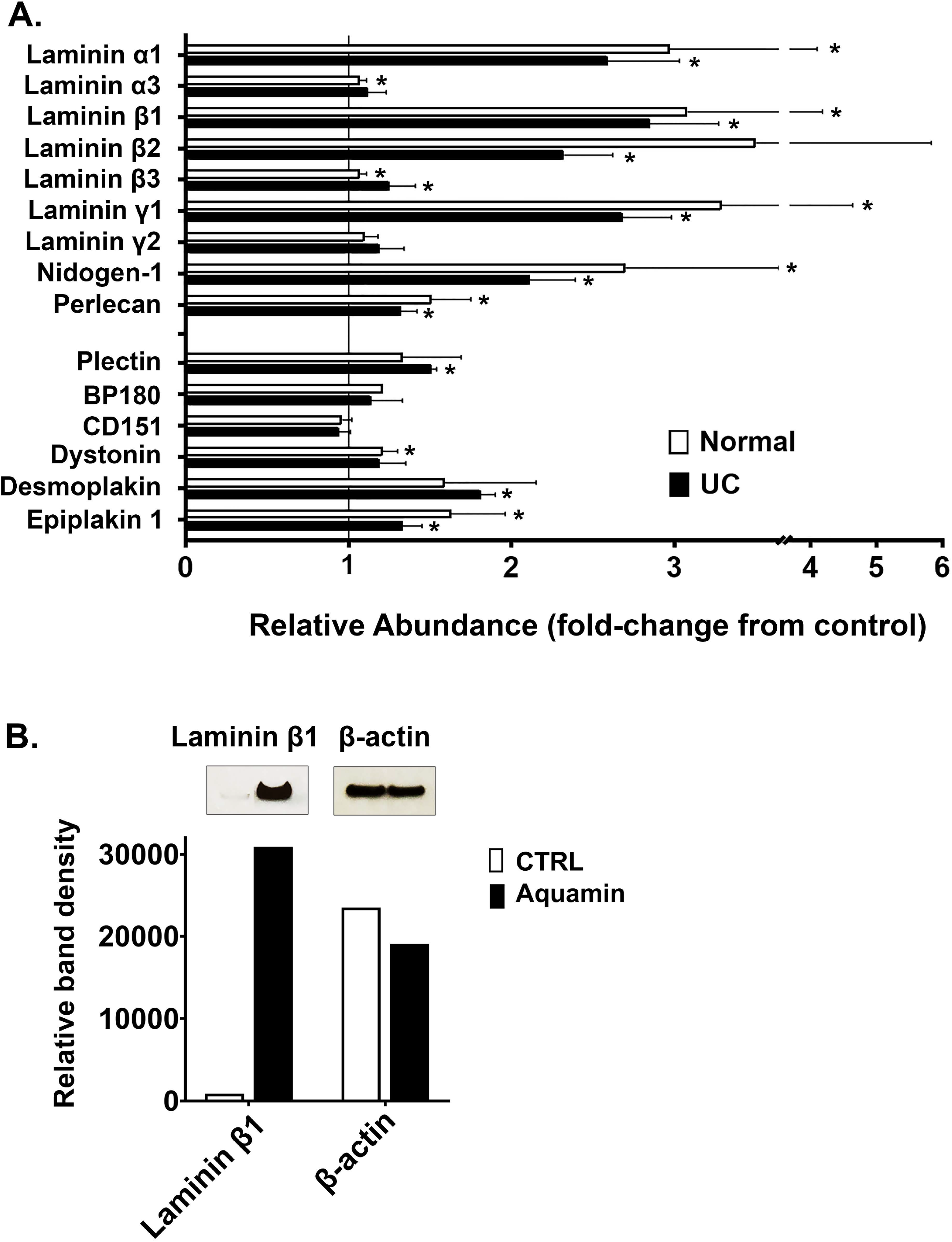
Effects of Aquamin^®^ on expression of proteins involved in cell-matrix interactions. **A**: Proteomic screen. Values reflect mean (±SD) fold-change from respective control in colonoids from normal subjects (N=4) and subjects with UC in remission (N=3). Methods for generating the data sets were described in detail in past reports (22,24). Data for individual proteins were compared for statistical differences using the Student t-test. Asterisk (*) indicates a difference from control at p<0.05. **B**: Western blotting. Protein isolated from each condition was assessed for laminin β1 expression by western blotting. 10 µg of protein from each condition was used. β-actin was assessed in parallel. Band quantitation was done using ImageJ software.

While this study did not address differentiation-related proteins, *per se*, we noted that keratin 8 and keratin 18 (components of intermediate filaments in gastrointestinal epithelial cells) (40) were increased in response to Aquamin^®^. With keratin 8, expression was increased 1.25 ± 0.01-fold and 1.52 ± 0.09-fold in the normal and UC data sets respectively. With keratin 18, values were 1.52 ± 0.41-fold and 1.86 ± 0.00-fold. Of interest, recent studies have demonstrated that mutations in Keratin 8/18 in colonic epithelial cells are associated with loss of permeability control in inflammatory bowel disease (41). Similarly, acute inflammation has been shown to reduce Keratin 8/18 expression; levels were restored with improvement in disease status as assessed by both clinical and endoscopic parameters (42).

In addition to the findings presented above, certain other proteins of interest were also detected in the protein screen. The α2 chain of type IV collagen was detected; it was modestly increased in response to Aquamin^®^. A number of integrin components were also detected, but components of the major laminin-binding integrins (α6, β1 and β4) were not significantly altered with Aquamin^®^ as compared to control. Among other moieties that have been reported to interact with laminin, both dystroglycan and syndecan were slightly down-regulated, sulfatide was unchanged and oncofetal antigen/immature laminin receptor OFA(iLRP)/67-kD laminin receptor was not detected. Finally, three proteins that serve as connectors between focal adhesions and the actin cytoskeleton (talin, vinculin and α-actinin) were detected but did not alter except vinculin, which was up-regulated in both sets (1.42- and 1.24-fold; in normal and UC respectively). Finally, we searched for the pathways associated with the proteins (presented in Fig 2A) using Reactome. Top 15 pathways with the involved entities are presented in Table 1. Type I hemidesmosome assembly, laminin interactions and extracellular matrix organization were at the top of the pathways list (Table 1).

**Table 1.** Top pathways associated with the proteins presented in Figure 2.

| *Pathway name* | *Submitted entities* |
| --- | --- |
| Type I hemidesmosome assembly | BP180;CD151;DST;LAMB3;LAMA3;LAMC2;PLEC |
| Laminin interactions | LAMB3;LAMA1;LAMB2;LAMA3;LAMB1;LAMC2;LAMC1;NID1;HSPG2 |
| Extracellular matrix organization | BP180;CD151;LAMB3;DST;LAMA1;LAMB2;LAMA3;LAMB1;LAMC2;LAMC1;NID1;HSPG2;PLEC |
| Non-integrin membrane-ECM interactions | LAMB3;LAMA1;LAMB2;LAMA3;LAMB1;LAMC2;LAMC1;HSPG2 |
| Degradation of the extracellular matrix | BP180;LAMB3;LAMB2;LAMA3;LAMB1;LAMC2;LAMC1;NID1;HSPG2 |
| Collagen formation | BP180;CD151;DST;LAMB3;LAMA3;LAMC2;PLEC |
| MET activates PTK2 signaling | LAMB3;LAMA1;LAMB2;LAMA3;LAMB1;LAMC2;LAMC1 |
| Assembly of collagen fibrils and other multimeric structures | BP180;CD151;DST;LAMB3;LAMA3;LAMC2;PLEC |
| MET promotes cell motility | LAMB3;LAMA1;LAMB2;LAMA3;LAMB1;LAMC2;LAMC1 |
| ECM proteoglycans | LAMA1;LAMB2;LAMA3;LAMB1;LAMC1;HSPG2 |
| Signaling by MET | LAMB3;LAMA1;LAMB2;LAMA3;LAMB1;LAMC2;LAMC1 |
| Cell junction organization | BP180;CD151;DST;LAMB3;LAMA3;LAMC2;PLEC |
| Cell-Cell communication | BP180;CD151;DST;LAMB3;LAMA3;LAMC2;PLEC |
| Anchoring fibril formation | LAMB3;LAMA3;LAMC2 |
| Signaling by Receptor Tyrosine Kinases | LAMB3;LAMA1;LAMB2;LAMA3;LAMB1;LAMC2;LAMC1 |

The pathway analysis report was generated by Reactome database (v75) for species “Homo sapiens.”

Western blotting with an antibody to the laminin β1 chain (most highly up-regulated of all the laminin chains detected in the proteomic screen) was used to confirm laminin up-regulation. Fig 2B shows the dramatic increase in laminin β1 expression in Aquamin^®^-treated organoids as compared to control.

### TEER values in organoids: Effects of Aquamin^®^ and anti-laminin treatment

Human colon organoids were plated on transwell membranes in growth medium. One day later, growth medium was replaced with the growth medium-KGM Gold mix (0.25 mM calcium; final concentration). In some wells, Aquamin^®^ was added to bring the final calcium level to 1.5 mM and provide the additional trace elements that make up the marine algae product. Electrical resistance across the cell layer was assessed as above. In the unsupplemented growth medium-KGM Gold mix, a TEER value of 1800 ohms per cm^2^ was achieved as compared to 2300 ohms per cm^2^ in differentiation medium (22% decrease) while the TEER value in Aquamin^®^ supplemented medium was virtually identical to that seen in differentiation medium (compare values in Fig 3 with those in Fig 1A).

**Fig 3.**
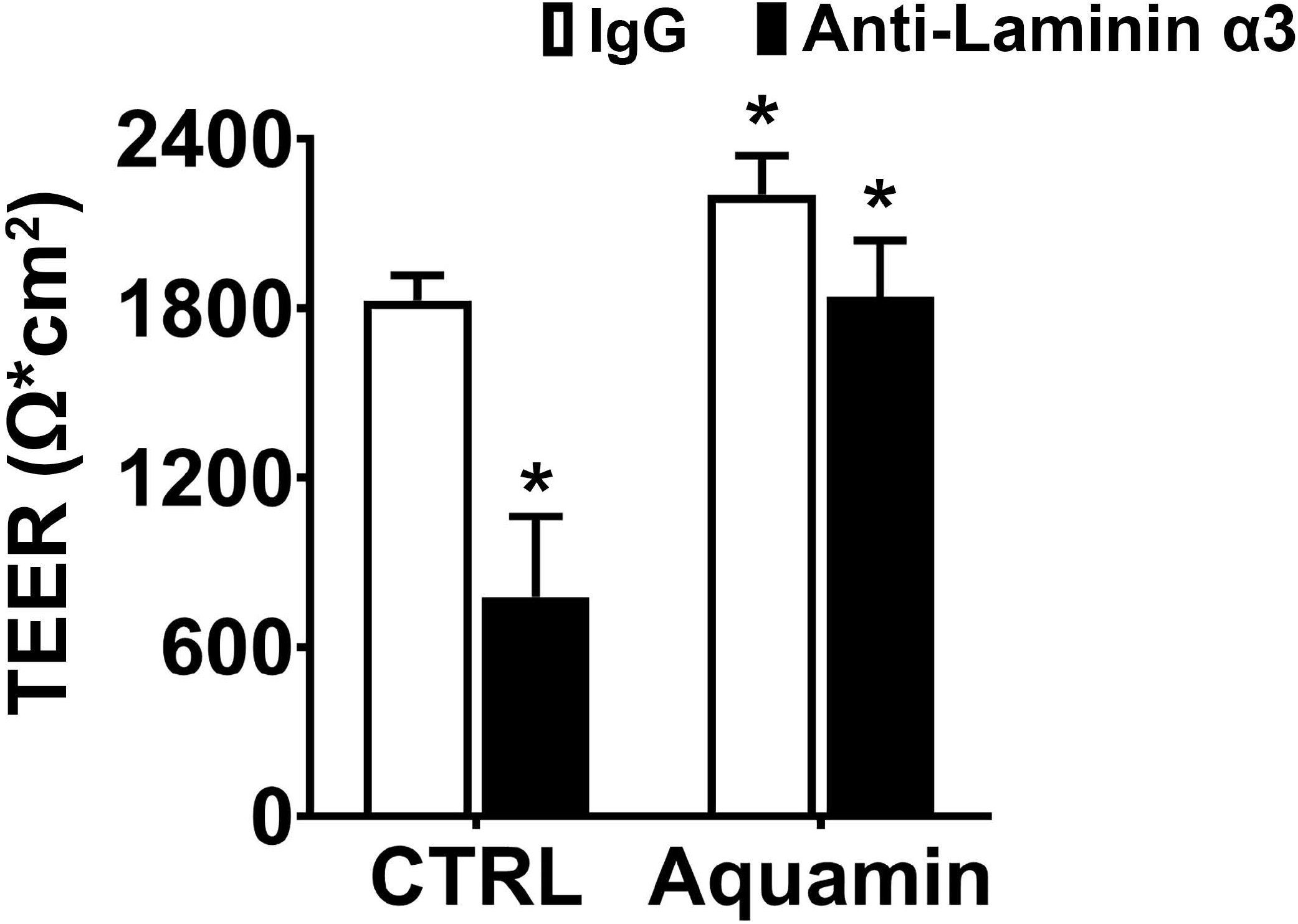
Transepithelial electrical resistance in growth medium-KGM Gold with or without Aquamin^®^ and with or without anti-laminin antibody. TEER values shown are means and standard deviations based on three separate experiments with 4 samples (individual membranes) per data point in each experiment. Effects of Aquamin^®^ and anti-laminin treatment on TEER were assessed. Data were compared for statistical differences using ANOVA followed by unpaired-group comparisons. Asterisk (*) above the open Aquamin^®^ bar indicates a difference from control at *p*<0.05. Asterisks (*) above the closed bars indicates difference from respective IgG control at p<0.05.

Fig 3 also shows the effects of anti-laminin treatment on TEER values in the two conditions. In Aquamin-supplemented medium, TEER values were reduced by 16% with anti-laminin. This is comparable to what was seen in differentiation medium (Fig 1B). In un-supplemented growth medium-KGM Gold, where TEER values were lower to begin, the inclusion of anti-laminin antibody further reduced TEER values by 57%.

### Effects of anti-laminin antibody on organoid cohesion

In our previous study, we demonstrated that treatment of human colon organoids with Aquamin^®^ increased organoid cohesion in parallel with TEER (23). To determine whether interactions involving laminin contributed to intra-organoid cohesion, organoids were maintained for a two-week period in growth medium-KGM Gold with and without anti-laminin. At the end of the incubation period, cohesion was assessed as described in Methods and in our previous study (23). As shown in Fig 4, no detectable antibody effect on organoid cohesion was seen. It was evident that there was no difference in the ratio of change in organoid size from post-to pre-harvest, the ratio was 0.45 and 0.46 with IgG and anti-laminin antibody, respectively (Fig 4).

**Fig 4.**
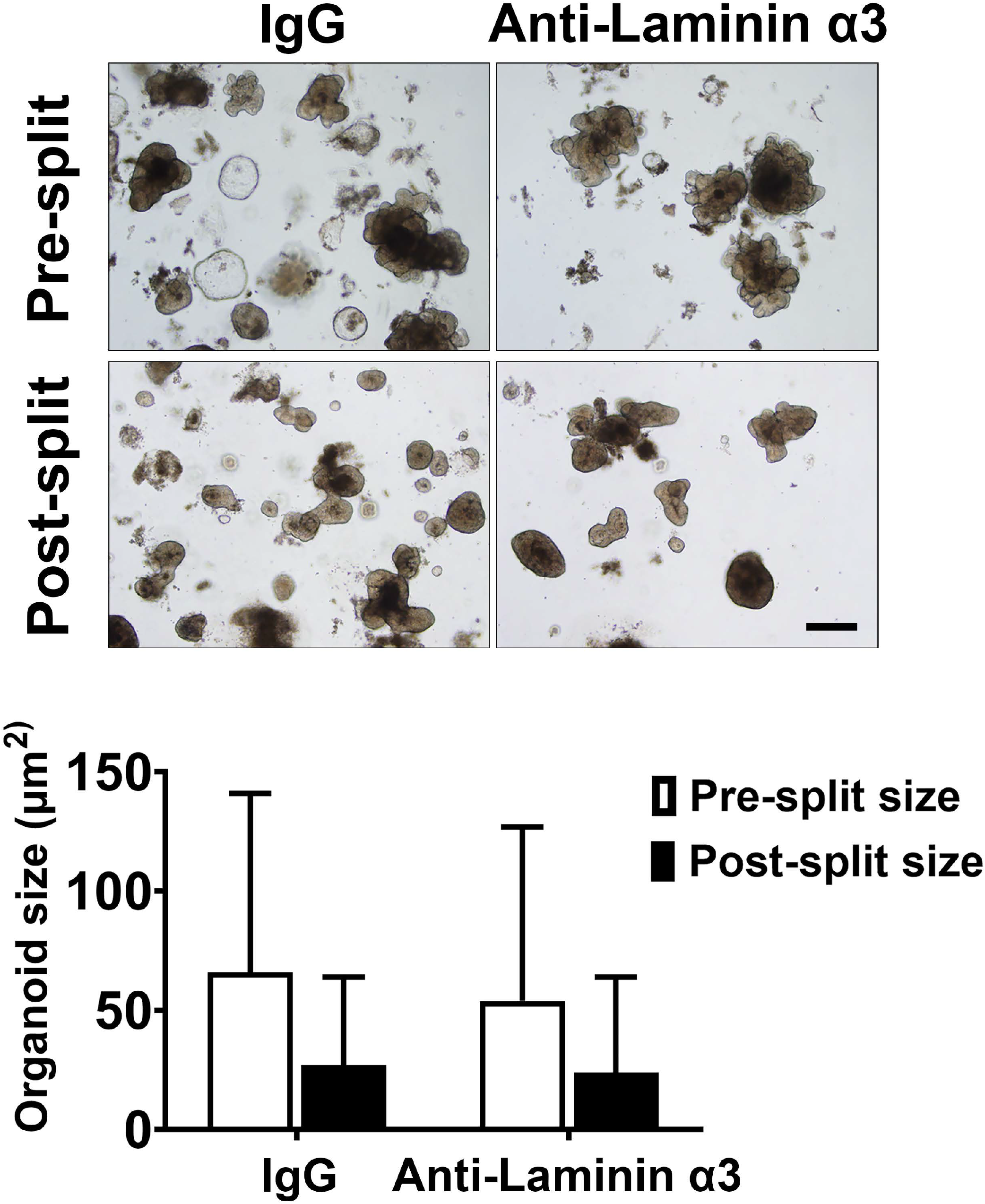
Colon organoid cohesion in growth medium-KGM Gold: Effect of anti-laminin antibody. Colon organoids were maintained for 14 days in growth medium-KGM Gold with and without anti-laminin. At the end of the incubation period, cell-cell cohesion was assessed. Values shown represent change in mean (±SD) surface area of individual colonoids based on two separate experiments with a minimum of 53 to 104 colonoids assessed individually per treatment group in both pre- and post-harvest cultures. Data were compared for statistical differences using ANOVA followed by unpaired-group comparisons. While the decrease in organoid size between post-harvest and pre-harvest groups were statistically significant with either IgG or anti-laminin, the differences between anti-laminin and IgG were not different. Inset: Representative examples of pre-harvest and post-harvest cultures. Scale bar = 200 µm

## DISCUSSION

Most studies of barrier dysfunction in the gastrointestinal tract have focused on the structural components that regulate cell-cell interactions (i.e., desmosomes and, especially, tight junctions) (16-19), but basement membrane disruptions are also commonly observed (25-28). Experimental animal models of colitis, likewise, demonstrate basement membrane disruptions in inflamed colonic tissue (28,43). In all of these settings, a loss or reduction in laminin immunoreactivity is commonly observed (25-28), although altered distribution of laminin forms has been reported as well, with some forms actually increasing (27). Laminin is not unique in being altered in inflammatory bowel conditions. Basement membrane collagens including both type IV and type V have been reported to be increased in inflamed bowel (28). Together, these past findings provide a picture of widespread cell-matrix disruption in the context of the inflamed colon. Although these changes are thought to be a consequence of the inflammatory process, anomalies have been noted in some patients with inflammatory bowel disease in the absence of acute tissue damage (4). Thus, pre-existing basement membrane irregularities may contribute to inflammation, and not simply be the consequence of tissue injury. In support of this, murine models in which laminin α-chain was over-expressed showed a decreased sensitivity to chemical-induced colitis (28). In another model, hemidesmosome disruption promoted colitis (43) in genetically manipulated animals.

Regardless of whether preexisting barrier defects in the gastrointestinal tract promote bowel inflammation or are simply the consequence of acute inflammation, improvement in barrier structure / function would seem to be of value. The findings presented here allow us to conclude i) that interfering with cell-basement membrane interactions reduces permeability control without a major effect on tissue cohesion in human colon organoid culture; ii) that treating colon organoids with a multi-mineral supplement leads to increased elaboration of basement membrane proteins and hemidesmosomal / intermediate filament components; and iii) that the increased elaboration of these proteins mitigates, at least partially, the consequences of interfering with cell-basement membrane interactions. As summarized graphically in the cartoon (Fig 5), the basement membrane, hemidesmosomal and intermediate filament proteins that are responsive to Aquamin^®^ treatment could be expected to have an effect on both focal adhesions and hemidesmosomes (44). In our previous studies, the same mineral supplement was shown to substantially increase desmosomes along the lateral surface of adjacent epithelial cells in colonoid culture without a major effect on tight junctions (22-24). Together with the data presented in our earlier study, the findings shown here suggest that while tight junctions maybe critical to permeability control in the gastrointestinal tract, both cell-cell and cell-matrix interactions are also important.

**Fig 5.**
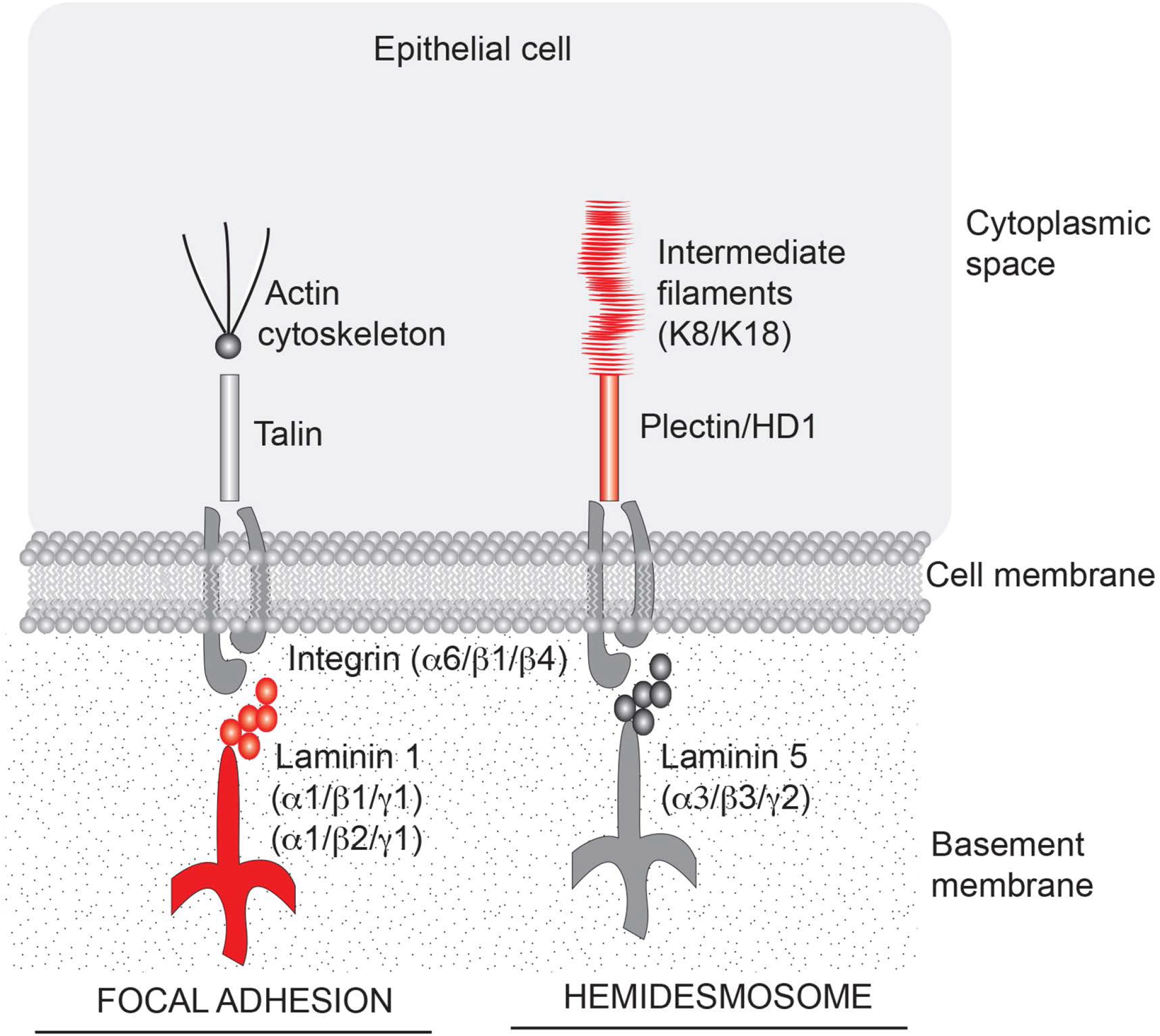
Aquamin^®^-responsive cell-matrix adhesion structures in the colon. Cartoon depicting structures important to cell-matrix adhesion in the colon and components of those structures that are responsive to Aquamin^®^ (shown in red). Based on the profile of proteins that are induced by Aquamin^®^, cell-matrix adhesion through both focal adhesions and hemidesmosomes could be affected.

The studies carried out here made use of a sophisticated *ex vivo* culture system, and whether the effects obtained *in vitro* have relevance to what occurs *in vivo* remains to be demonstrated. In an effort to begin addressing this issue, we have recently carried out a pilot phase trial in which ten healthy subjects were treated with the same multi-mineral product (Aquamin^®^) used here. To summarize the results of this pilot study, there were no tolerability issues with daily Aquamin^®^ ingestion over a 90-day period and no safety concerns (45). Equally important, when Aquamin^®^-treated subjects were compared to subjects receiving placebo for the same period, we saw up-regulation of laminin chains along with increased levels of other basement membrane components and hemidesmosome moieties in colonic biopsies (46).

Subjects receiving calcium alone (i.e., the most abundant mineral in Aquamin^®^) also demonstrated increases in several of the same molecules, but the degree of up-regulation with calcium alone was lower than that seen with Aquamin^®^ (46).

As a follow-up, we are conducting a 180-day interventional trial with Aquamin^®^ in UC patients (ClinicalTrials.gov: NCT03869905). In addition to evaluating therapeutic benefit, the same approaches used in the earlier trial with healthy individuals (immunohistology and proteomics) are being used to evaluate proteins changes in the colon over the course of intervention. In parallel, the urine lactulose/mannitol ratio (47) is being assessed to provide a direct measure of treatment effects on gastrointestinal permeability (ClinicalTrials.gov: NCT04855799). If successful, Aquamin^®^ or a similarly formulated product could be used as a low-cost, low-to no-toxicity adjuvant therapy to improve gastrointestinal barrier function in individuals suffering from a variety of gastrointestinal maladies. At the very least, individuals with barrier defect-associated gastrointestinal conditions should be encouraged to include an adequate source of calcium and other minerals in their diet. Unfortunately, deficiencies in calcium and other critical mineral components are widespread throughout the world (48) and this is especially true for those consuming a Western-style diet (49).

Finally, there is another group of diseases – epidermolysis bullosa and related conditions – that are manifestations of mutations in various basement membrane, desmosomal / hemidesmosomal and keratin genes (50). At the same time, there are case reports and studies that provide evidence of an association between bullous pemphigoid and inflammatory bowel disease (51-53). At this point, we can only speculate as to whether optimizing the expression of multiple cell-cell and cell-matrix adhesion molecules in an individual might overcome, at least in part, the consequences of a function-modifying mutation in one or another critical component. If this turns out to be the case, it could open the door to a new adjuvant therapeutic approach. While speculative for now, experimental models in which a hypothesis could be tested are available (54-56).

In summary, an intact permeability barrier is required for healthy gastrointestinal function. While cell-cell adhesion structures are well-known participants in effective permeability control, the present study provides evidence that cell-matrix interactions are also important. These studies show, furthermore, that a multi-mineral natural product has the capacity to stimulate the production of cell-matrix adhesion moieties and, concomitantly, to improve barrier control.

## DATA AVAILABILITY

All relevant data are within the manuscript and its Supporting Information files. The mass spectrometry proteomics data are available on ProteomeXchange Consortium (PRIDE partner repository) – identifier PXD020244.

## ACKNOWLEDGMENTS

We thank Marigot LTD (Cork, Ireland) for providing Aquamin^®^ as a gift. We thank the Microscopy and Imaging Laboratory (MIL) for help with confocal fluorescence microscopy. We thank the Translational Tissue Modeling Laboratory (TTML) at the University of Michigan for help with colon organoid propagation and help with the TEER assay. We also thank the Proteomics Resource Facility (Pathology Department) for help with proteomic data acquisition.

